# Olfactory bulb-medial prefrontal cortex circuit slow oscillations encode working memory representations

**DOI:** 10.1101/2025.10.30.685257

**Authors:** Ali Samii Moghaddam, Morteza Mooziri, Mohammad Reza Raoufy, Mohammad Ali Mirshekar

**Affiliations:** Genetics of Non-Communicable Disease Research Center, Zahedan University of Medical Sciences, Zahedan, Iran; Department of Physiology, Faculty of Medical Sciences, Tarbiat Modares University, Tehran, Iran; Institute for Brain and Cognition, Faculty of Medical Sciences, Tarbiat Modares University, Tehran, Iran; Clinical Immunology Research Center, Zahedan University of Medical Sciences, Zahedan, Iran; Department of Physiology, School of Medicine, Zahedan University of Medical Sciences, Zahedan, Iran

**Keywords:** Working memory, Medial prefrontal cortex, Olfactory bulb, Brain oscillations

## Abstract

Working memory (WM), the ability to maintain task-related information, is fundamental to animal behaviors. Despite significance, there is limited knowledge on how it is orchestrated by the neural activity in the brain. Here, we show that the olfactory bulb (OB) and medial prefrontal cortex (mPFC) slow oscillations (1-30 Hz) are augmented during WM, and furthermore, the pattern of these activities convey task-related information. Moreover, we find that the OB-mPFC delta and beta power-based connectivity is enhanced by WM. On the other hand, theta oscillations have the most prominent roles in phase coupling in this circuit. We also show bidirectional information transfer, stronger in the mPFC-to-OB direction, between the two brain areas. Furthermore, during WM, beta activity in both regions is tuned by the phase of local and long-range delta as well as theta oscillations. Together, our results suggest that the dynamics of OB-mPFC circuit slow band activities underlies WM, with possible implications for other cognitive functions in health and disease.

## Introduction

Working memory (WM), the ability to maintain task-related information for immediate use, is fundamental to mammalian cognition^1-4^. Importantly, WM is an essential building block for a wide range of complex goal-directed behaviors, from learning to problem-solving and others^1-4^. Furthermore, WM is disrupted in several neuropsychiatric disorders, such as schizophrenia and attention-deficit/hyperactivity disorder (ADHD)^5-7^. However, there is limited knowledge on its underlying neural architectures.

There is substantial evidence that processing of WM requires coordinated activity of a network of brain regions^1-4,8-12^. In this line, prefrontal cortex (PFC), as a key neural hub for the expression of a wide range of emotional and cognitive behaviors, is believed to have the most prominent contributions^1,3,4,8-10,12-19^. Importantly, presence of long-range functional and/or structural connections between PFC and other brain areas, such as the medial temporal lobe (MTL), furthers signifies its critical roles to form behaviors^1,4,9,10,13-25^. Specifically, a large body of human^21,22,26^, non-human primate (NHP)^1,3,4,8,20,27-31^, and rodent^9,10,25,32^ studies have shown that PFC slow oscillations support WM memory through diverse regional and inter-regional mechanisms. Therefore, local activities and long-range effects of PFC are pivotal for keeping the information in WM.

Previous studies show that olfactory bulb (OB) participates in different cognitive and emotional behaviors, including WM^10,11,17-19,32-34^, which happens through local processing and extensive connections to several brain regions, including medial PFC (mPFC) and MTL^10,11,14,17-19,23,32-38^. Removing or lesioning OB disrupts regional and inter-regional brain oscillations and impairs cognition in areas like attention as well as short- and long-term memory^14,39-43^. Also, clinically important, OB dysfunction is suggested to be associated with neurocognitive and psychiatric disorders^10,11,14,17-19,33,36,43,44^. For instance, olfactory impairment signals the emergence of prodromal Alzheimer disease^44^. Furthermore, conditions associated with respiratory dysfunction often induce abnormalities in the activity of brain regions and disturb behavior^9,14,17,19,45-47^. Therefore, OB is a major brain structure to support appropriate expression of different behaviors.

Brain oscillations are important means to coordinate the activity of brain regions and are often behaviorally relevant^4,9-11,13-19,24,25,32,33,45,47,48^. Notably, slow oscillation are prominent brain rhythms to mediate a wide range of communications between distant structures^4,10,11,13-19,32,33,47^. Importantly in this line, there are anatomical and functional connectivity between OB and mPFC^10,11,14,17,18,23,33^, whose enhancements are associated with behaviors like WM, anxiety, and fear^10,13,14,17,18^. Furthermore, expression of these behaviors often accompanies bidirectional information transfer between the two regions^14,17,18^.

Therefore, together, PFC and OB underlie a wide range of animal behaviors^1,3,4,8-11,13-19,33,39,45,47,49^. While a vast body of literature has focused on the roles of mPFC slow oscillations, there is little knowledge on how OB support WM in the mammalian brain. Here, we used behavioral and neurophysiological techniques to study the dynamics of slow band (1-30 Hz) neural activity in the rodent OB-mPFC circuit during WM.

Real-time simultaneous recordings from mPFC and OB revealed that WM boosts mPFC and OB slow oscillations in a manner that the pattern of these neural activities contains information on the animal’s experience of a cognitive process. Moreover, we found that during WM, the OB-mPFC circuit functional connectivity is enhanced as the following: first, we observed enhanced power-based connectivity, which happens through delta (1-4 Hz) and beta (12-30 Hz) bands; on the other hand, while nearly all frequencies show greater phase coupling during WM, this effect is most prominent in theta (4-12 Hz) oscillations. Also, we report bidirectional slow band information transfer between mPFC and OB. Lastly, similar to primates^8^, we show that the phase of delta and theta oscillations modulate the regional and distant beta activity to support the maintenance of information in WM.

## Results

### Neurophysiology and behavior

We recorded the mPFC and OB LFPs, while the rats were freely exploring a Y-maze apparatus (Fig. 1), which is a common procedure for evaluating rodent short-term memory, following a baseline recording session. A correct trial in this setting is defined once the animal visits all three arms consecutively (all other possibilities are considered as wrong trials; Fig. 1B).

**Figure 1.**
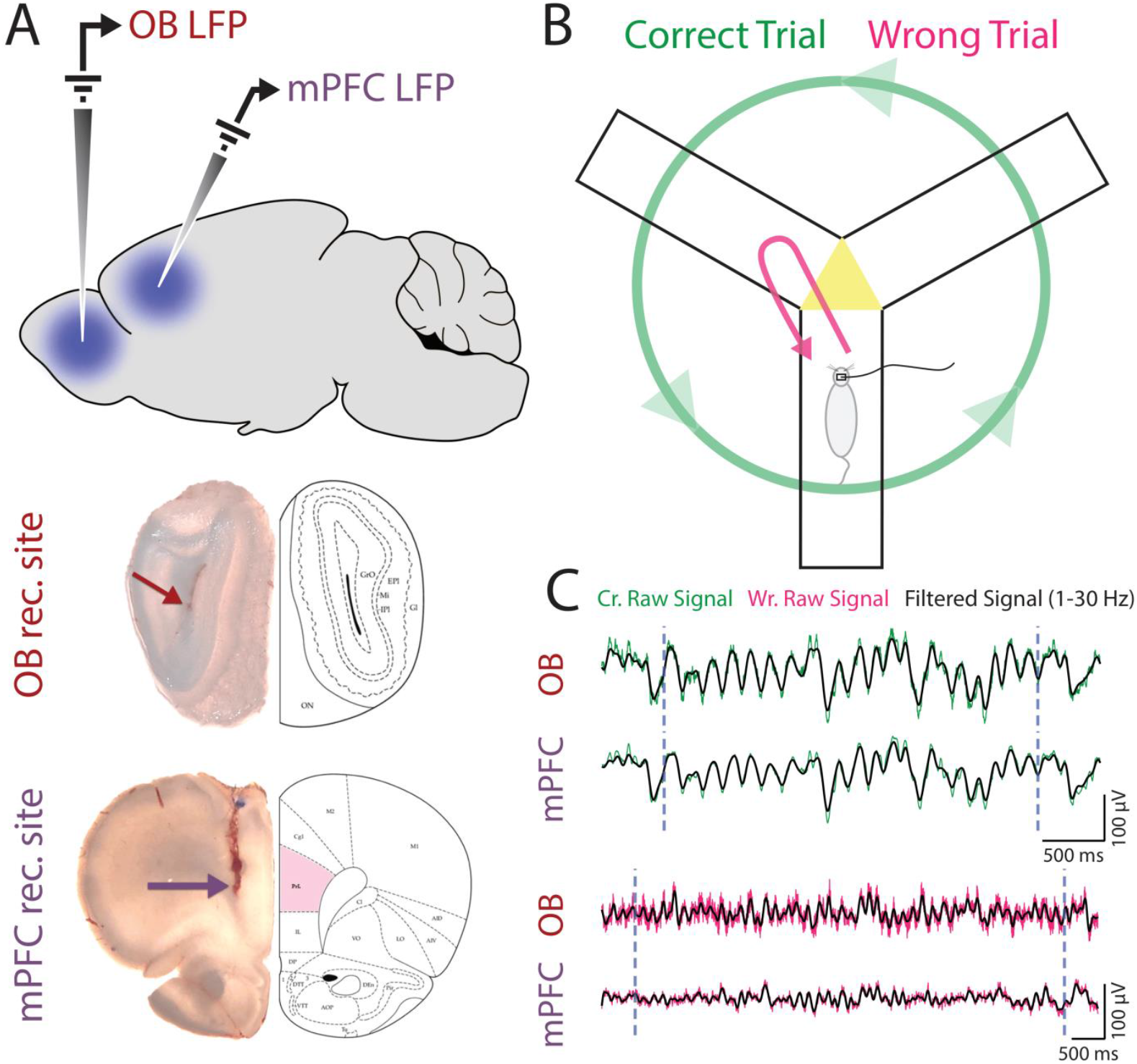
Experiment. (A) Schematic (upper panel) and histological verification of OB (middle panel) and mPFC (lower panel) electrode sites. (B) Schematic of the behavioral task, i.e., Y-maze. Green/pink trajectory denotes a correct/wrong trial. (C) Sample OB and mPFC LFP signal during a correct (green) and wrong (pink) transition. Vertical grey dashed lines indicate the start (entering the central triangle) and end (leaving the central triangle) of the trial. LFP, local field potential; OB, olfactory bulb; mPFC, medial prefrontal cortex.

### WM execution enhances the mPFC and OB slow oscillations

Low-frequency neural activity in various brain regions and among different species are shown to underlie cognitive and emotional processes, including short-term maintenance of information^3,13-15,17,18,31,32,45,47,49^. To study the mPFC and OB slow oscillations during rodent spatial working memory, we evaluated each region’s power spectral density (PSD) at delta, theta, and beta frequency bands. As shown Fig. 2A, WM execution enhances the mPFC slow oscillations, regardless of performance (frequencies that change compared to the baseline are denoted with horizontal green/pink line for correct/wrong trials in Fig. 2A; thin/thick line shows statistical significance at the p-value of 0.05/0.001 from permutation testing). As expected, mPFC delta (mean ± SEM; correct = 6.74 ± 2.03, wrong = 2.41 ± 0.80, p = 0.048; Fig. 2A,B) and beta (mean ± SEM; correct = 6.06 ± 2.06, wrong = 1.85 ± 0.88, p = 0.058; Fig. 2A,B) enhancements are greater during correct trials, compared to wrong trials; however, mPFC theta activity did not differ between the two conditions (mean ± SEM; correct = 11.08 ± 3.53, wrong = 4.40 ± 2.38, p = 0.12; Fig. 2A,B). Additionally, we observed that WM task-involvement also boosts OB slow activity (Fig. 2C; statistical significance denoted in horizontal green and pink lines and should be interpreted as in Fig. 2A). Further, OB delta (mean ± SEM; correct = 3.23 ± 0.98, wrong = 0.75 ± 0.39, p = 0.02; Fig. 2C,D) and beta (mean ± SEM; correct = 1.94 ± 0.72, wrong = 0.70 ± 0.19, p = 0.03; Fig. 2C,D) enhancements are greater during correct trials, compared to wrong trials; this effect, despite presence of the trend, did not reach significance in theta band (mean ± SEM; correct = 5.92 ± 1.08, wrong = 3.67 ± 0.78, p = 0.095; Fig. 2C,D).

**Figure 2.**
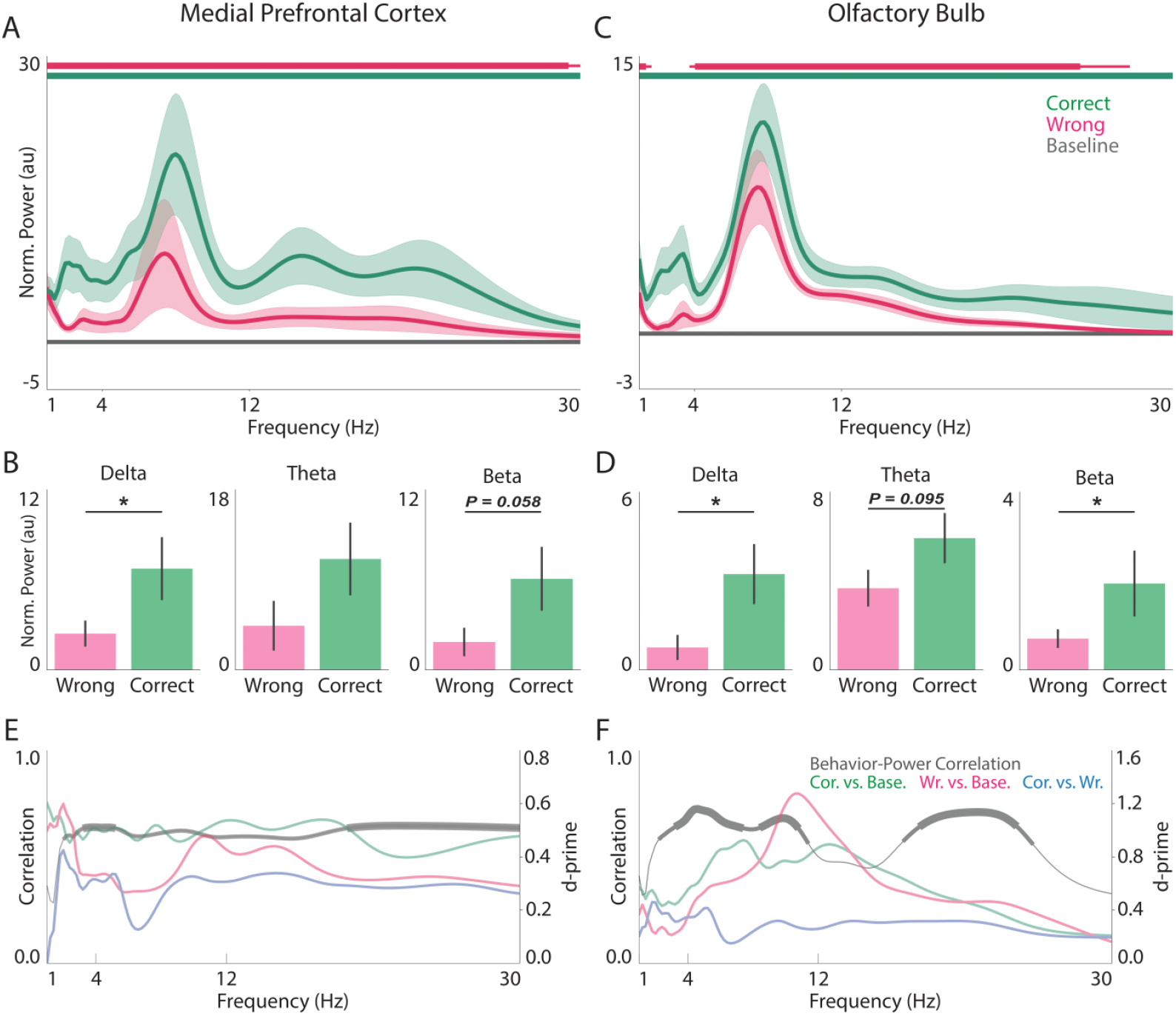
mPFC and OB regional slow oscillations during WM. (A,C) Normalized PSD of mPFC (A) and OB (B) during baseline, wrong, and correct trials. Solid line and shaded area denote mean and SEM of power, respectively. Horizontal green/pink line indicates frequencies with statistical difference between correct/wrong and baseline trials. Thin/thick line represents statistical significance at the p-value level of 0.05/0.001. (B,D) Statistical comparison of average delta (leftmost panels), theta (middle panels), and beta (rightmost panels) power of mPFC (B) and OB (D) between correct and wrong trials. Bars and error bars represent mean and SEM, respectively. (E,F) Frequency-wise power-behavior correlation (left y-axes) and magnitude of power difference i.e., d’, between conditions (right y-axes) for mPFC (E) and OB (F). Thin correlation line indicates values with p ≥ 0.1, while medium/thick line indicates correlation values with p < 0.1/0.05. Permutation test. * p < 0.05. OB, olfactory bulb; mPFC, medial prefrontal cortex; PSD, power spectral density.

To measure the strength of activation, we used d-prime (d’; see Methods) to compare different conditions in each region and each frequency. In mPFC, when compared to baseline activity, correct trials had a medium effect size in all studied frequencies (average d’ = 0.48, 0.40 < d’ < 0.60; green trace in Fig. 2E); while wrong trials were generally comparable to corrects in the delta band (average d’ = 0.45, 0.33 < d’ < 0.60), they had smaller effects in both theta (average d’ = 0.35, 0.27 < d’ < 0.48) and beta (average d’ = 0.35, 0.29 < d’ < 0.44) frequencies (red trace in Fig. 2E). Moreover, when directly comparing correct vs. wrong, the difference was more prominent in delta (average d’ = 0.28, −0.01 < d’ < 0.43), higher theta (8-12 Hz; average d’ = 0.31, 0.26 < d’ < 0.33), and beta (average d’ = 0.30, 0.26 < d’ < 0.34) bands, with a pronounced wide-band drop in lower theta (4-8 Hz; average d’ = 0.23, 0.13 < d’ < 0.34) frequencies (blue trace in Fig. 2E). In the OB, we generally observed larger effects, i.e., stronger WM-related activations. Specifically, in correct trials, the difference with baseline was greater in theta band (average d’ = 0.79, 0.58 < d’ < 0.93), with delta (average d’ = 0.49, 0.43 < d’ < 0.57) and beta (average d’ = 0.48, 0.21 < d’ < 0.89) oscillations showing slightly weaker activations (green trace in Fig. 2F); also, a similar trend was observed in the wrong trials (delta: average d’ = 0.28, 0.21 < d’ < 0.44; theta: average d’ = 0.84, 0.42 < d’ < 1.28; beta: average d’ = 0.49, 0.16 < d’ < 1.09; red trace in Fig. 2F). Further, the correct vs. wrong difference followed the general trend observed in mPFC (delta: average d’ = 0.35, 0.21 < d’ < 0.46; low theta: average d’ = 0.27, 0.15 < d’ < 0.42; high theta: average d’ = 0.28, 0.26 < d’ < 0.32; beta: average d’ = 0.27, 0.19 < d’ < 0.32; red trace in Fig. 2F).

The above results suggest functional heterogeneities in different frequencies of theta oscillations. Similarly, hippocampal theta sub-bands are specialized for different purposes, with slower/faster oscillations supporting emotional/cognitive processing^50,51^. In this line, we have formerly shown that during anxiety, both mPFC and OB low theta enhancements are more pronounced, compared to high theta^18^, which is supported by further rodent and human evidence on slow theta processing of emotions in regions like amygdala, mPFC, and OB^14-17^. By dividing theta oscillations into low and high sub-bands, as defined above, we observed that low theta activations did not significantly differ between correct and wrong trials in neither region (mean ± SEM; mPFC: correct = 11.90 ± 3.87, wrong = 5.53 ± 3.33, p = 0.23; OB: correct = 6.55 ± 1.23, wrong = 4.23 ± 1.11, p = 0.17; Fig. 2A,B). Interestingly, high theta activity was greater in correct, compared to wrong trials, in both mPFC (correct = 11.90 ± 3.87, wrong = 3.28 ± 1.47, p = 0.066; Fig. 2A) and OB (correct = 5.29 ± 1.07, wrong = 3.11 ± 0.50, p = 0.069; Fig. 2B). Note that these differences are related to WM content, and do not represent task-involvement, in which case the comparisons should be made against the baseline activities. Importantly, this data suggest that low theta frequencies have a role in detection of a cognitive demand (task vs. baseline in Fig. 1A,B,E,F).

Next, we were curious to see if the regional neural activity is correlated with behavior. We observed positive correlations between mPFC power and the animals’ performance in a wide-band manner; lower delta oscillations, which have the smallest correct-wrong d’, remained spared here too (grey trace in Fig. 2E). Also, OB activity was correlated with the behavior in multiple frequencies, primarily spanning the higher delta, which is similar to mPFC-behavior correlation and is associated with relatively larger (compared to lower delta) correct-wrong d’ in regional activity, as well as theta and mid-beta bands (grey trace in Fig. 2E). Overall, these results show that regional slow oscillations of the mPFC and OB are augmented in a WM process for the following purposes: 1) that the animal detects a situation in which WM, as a cognitive process, is required to guide through the environment (task vs. baseline); and 2) storage of information for an immediate use (correct vs. wrong).

### mPFC and OB slow oscillations encode WM representations

Pattern of activities in a population of neurons define sensory and/or cognitive representations^52-56^ (Fig. 3A, left panel). If the same notion holds true in a hypothetical neural space where each dimension is the activity of a given region in a frequency band (Fig. 3A, right panel), then task-related information might potentially be readable from such a space. Thus, we tried to categorize different task features from pattern of oscillatory activities, using a linear classifier (see Methods). While the neural activity of neither region conveyed notable trial type information, i.e., correct vs. wrong, (mean ± SD; mPFC: accuracy_observation_ = 49.80 ± 5.15, accuracy_shuffled_ = 49.18 ± 5.54, p = 0.25; OB: accuracy_observation_ = 51.38 ± 6.09, accuracy_shuffled_ = 49.30 ± 7.25, p = 0.003; permutation test; data not shown), we were able to categorize task vs. resting state with both mPFC (mean ± SD; accuracy_observation_ = 63.62 ± 8.24, accuracy_shuffled_ = 49.71 ± 3.07, p < 1e-4; permutation test; Fig. 3B) and OB (mean ± SD; accuracy_observation_ = 84.49 ± 3.91, accuracy_shuffled_ = 49.56 ± 4.84, p < 1e-4; permutation test; Fig. 3B) activity; interestingly, in the later setting, OB slow oscillations conveyed more robust information to distinguish a condition with high cognitive load, compared to mPFC (d’_mPFC_ = 2.37, d’_OB_ = 7.94).

**Figure 3.**
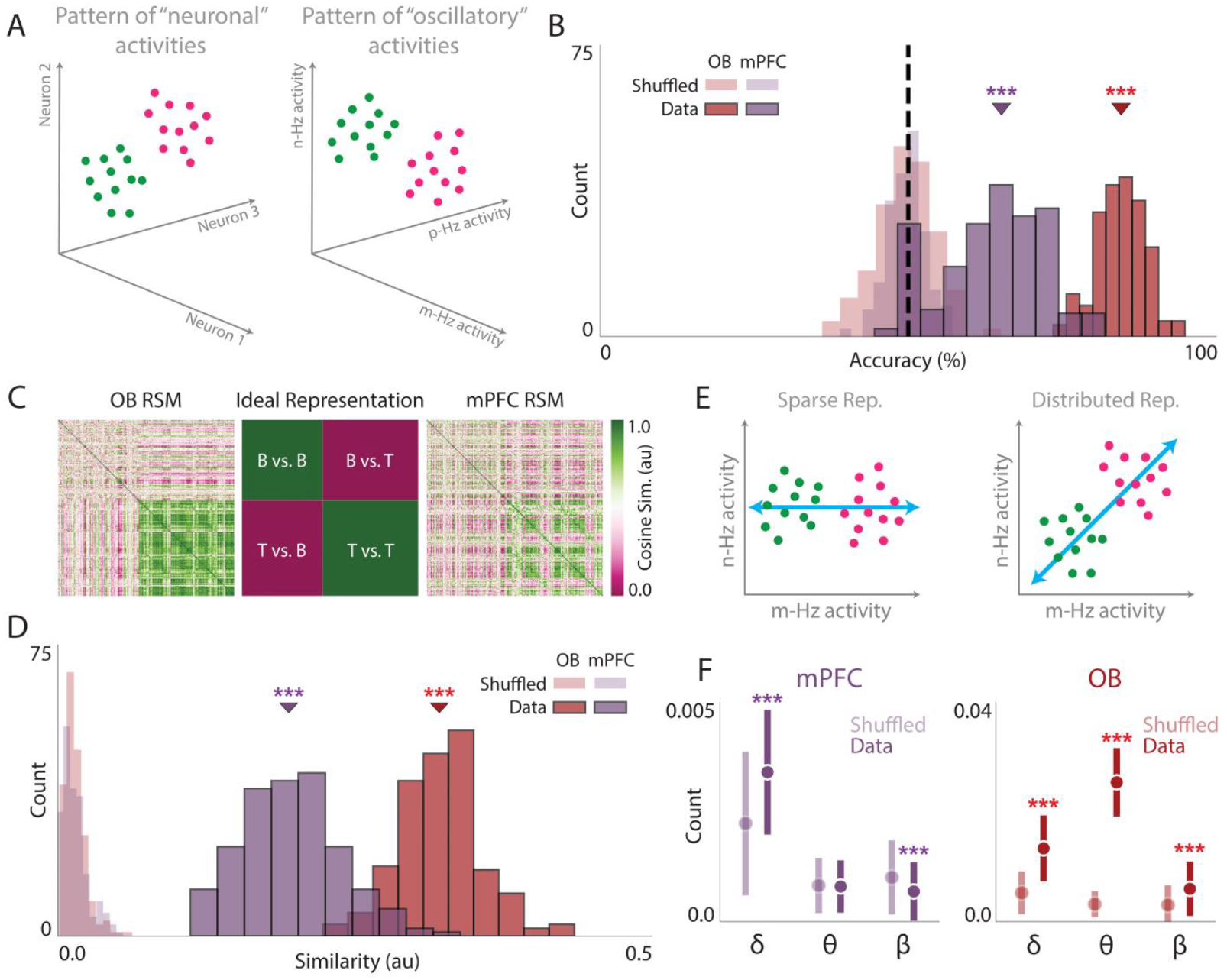
WM representation in OB and PFC slow oscillations activity pattern. (A) Schematic illustration of the hypothetical high-dimensional neural space when the features are neurons (left panel) or oscillatory activities (right panel). (B) Histogram of the linear classifiers’ performance trained to detect baseline vs. task from the pattern of mPFC and OB slow oscillations. Arrowheads denote the median of the performance distributions. (C) RSMs of OB (leftmost panel) and mPFC (rightmost panel) slow oscillation activities as well as the ideal representation (middle panel). (D) Histogram of the similarity between data-derived and ideal RSMs for the task-baseline setting. Arrowheads denote the median of the similarity distributions. (E) Schematic representation of sparse (left panel) vs. distributed (right panel) representation in a high-dimensional space. Blue line indicates the encoding axis in the hypothetical space. (F) Statistical comparison of the classifiers’ weights, trained on task vs. baseline, between normal and label-shuffled classification for each frequency band in the mPFC (left panel) and OB (right panel). Point and error bars represent mean and SD, respectively. Permutation test. *** p < 0.001. OB, olfactory bulb; mPFC, medial prefrontal cortex; RSM, representational similarity matrix.

To make more concrete evaluations, we tried to use representational similarity analysis (RSA; see Methods)^52^. For that, initially, we created ideal representational similarity matrices (RSMs) for correct vs. wrong (RSM_WC_) and task vs. baseline (RSM_BT_) settings. Next, data-derived RSMs were formed from the neural activity of either region for each condition, and were compared to the corresponding ideal representations (Fig. 3C; see Methods). Both mPFC (mean ± SD; similarity_observation_ = 0.02 ± 0.01, similarity_shuffled_ = 0.01 ± 0.01, p < 1e-4; permutation test; data not shown) and OB (mean ± SD; similarity_observation_ = 0.015 ± 0.005, similarity_shuffled_ = 0.013 ± 0.01, p = 0.01; permutation test; data not shown) had weak correct-wrong distinctions, with mPFC forming slightly more distinctive representations (d’_mPFC_ = 1.12, d’_OB_ = 0.25). Also, both regions formed strong task vs. baseline representations (mean ± SD; mPFC: similarity_observation_ = 0.20 ± 0.04, similarity_shuffled_ = 0.01 ± 0.01, p < 1e-4, d’ = 6.05; OB: similarity_observation_ = 0.32 ± 0.03, similarity_shuffled_ = 0.01 ± 0.01, p < 1e-4, d’ = 12.18; Fig. 3C,D). Logically, RSA considers patterns that provide meaningful information, which is especially the case in sensory systems where different categories or classes of data require specific processing circuitries. However, in the current setting, baseline activities are more probably spontaneous and without an underlying population processing. By visual inspection, this pattern is obvious in both mPFC and OB representations (Fig. 3C). Therefore, we also compared the data-derived RSMs against an ideal representation that does not consider the baseline neural activity to be similar in different occasions (Supplementary Fig. 1A), and found stronger OB (mean ± SD; similarity_observation_ = 0.48 ± 0.03, similarity_shuffled_ = 0.02 ± 0.04, p < 1e-4; d’ = 12.93; permutation test; Supplementary Fig. 1B), however not mPFC (mean ± SD; similarity_observation_ = 0.18 ± 0.04, similarity_shuffled_ = 0.01 ± 0.02, p < 1e-4; d’ = 5.26; permutation test; Supplementary Fig. 1B), similarities. Overall, these results show that beyond enhancement, the pattern of mPFC and OB slow oscillations encode WM information.

Next, we asked what kind of neural architecture could potentially underlie these processes. Specifically, whether either region forms sparse or distributed WM representations; in case of a sparse neural representation, a few dimensions in the high-dimensional neural space define most of the separation observed between classes of data (Fig. 3E, left panel); on the other hand, once several dimensions contribute to a separation, the resulting representations are defined as distributed (Fig. 3E, right panel). To this aim, we looked at the weights extracted from our LDA classifier for the task-baseline condition (see Methods and Fig. 3B). In the mPFC, delta band weights were greater in data, compared to shuffled classification (mean ± SD; weight_observation_ = 3e-3 ± 1e-3, weight_shuffled_ = 2e-3 ± 2e-3, p < 1e-4; permutation test; Fig. 3F, left panel); this effect did not exist in neither theta (mean ± SD; weight_observation_ = 1e-3 ± 1e-3, weight_shuffled_ = 1e-3 ± 1e-3, p = 0.70; permutation test; Fig. 3F, left panel) or beta (mean ± SD; weight_observation_ = 7e-4 ± 7e-4, weight_shuffled_ = 10e-4 ± 8e-4, p < 1e-4; permutation test; Fig. 3F, left panel) bands. In the OB, all frequency bands showed greater weights in the data, compared to the shuffled (mean ± SD; delta: weight_observation_ = 13e-3 ± 6e-3, weight_shuffled_ = 5e-3 ± 4e-3, p < 1e-4; theta: weight_observation_ = 25e-3 ± 6e-3, weight_shuffled_ = 3e-3 ± 2e-3, p < 1e-4; beta: weight_observation_ = 6e-3 ± 5e-3, weight_shuffled_ = 3e-3 ± 4e-3, p < 1e-4; permutation test; Fig. 3F, right panel), with the theta oscillations having strongest contributions (d’_delta_ = 1.60, d’_theta_ = 4.71, d’_beta_ = 0.68). Therefore, while OB has a distributed WM representation, delta oscillations are the prominent contributors in defining the mPFC readout.

### OB-mPFC slow band functional connectivity is enhanced during WM

Low frequency oscillations synchronize the neural activity across brain regions for different cognitive purposes, including WM^9-11,13-19,33,45,47^. Thus, we used power correlation to evaluate the OB-mPFC coupling (Fig. 4A). During baseline, we observed weak connectivity in the delta, theta and mid-beta bands (Fig. 4B,C). The OB-mPFC connectivity in wrong trials was predominantly present and enhanced, compared to baseline, in delta and low-beta oscillations (Fig. 4B-D). Besides, we observed wide-band coupling during the correct trials, which was enhanced in delta and beta frequencies, compared to baseline (Fig. 4B-D). While theta power correlation existed in both correct and wrong conditions, it was weaker, compared to delta and low-beta, and was not significantly enhanced in neither condition compared to baseline (Fig. 4B-D). Moreover, correct trials had greater theta and beta, but not delta, connectivity compared to wrong trials (Fig. 4C,D). Overall, these results suggest that delta and beta bands are the prominent oscillations to mediate OB-mPFC power-based connectivity during WM.

**Figure 4.**
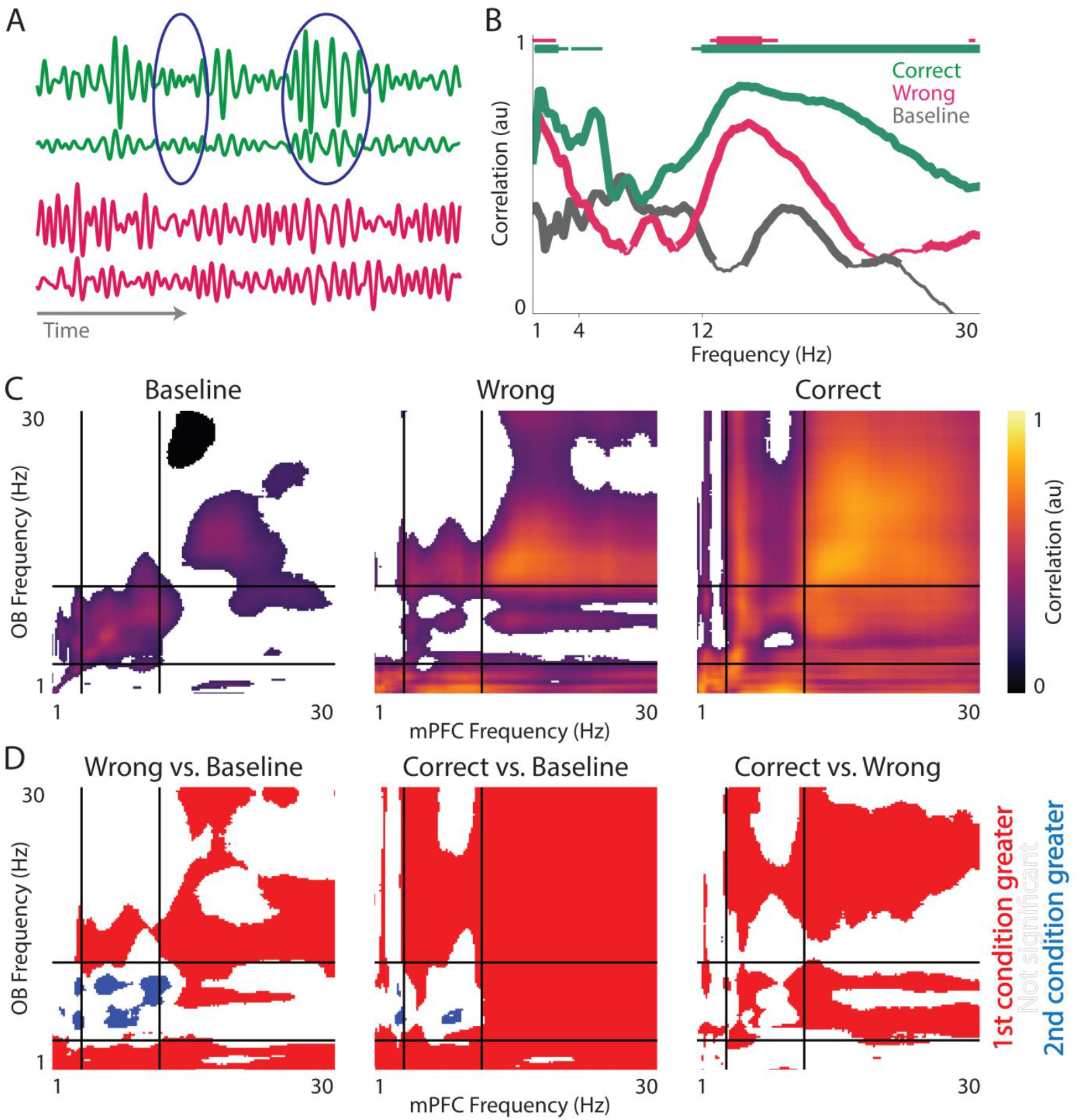
OB-mPFC power-based connectivity across slow oscillations during WM. (A) Schematic of the relationship between the power of two signals in the presence (green) or absence (pink) of power-correlation. (B) Frequency-wise power-correlation between OB and PFC in baseline, wrong and correct conditions. Thick/thin line represents significant/insignificant correlation values at p = 0.05. Horizontal green/pink line denotes frequencies in which power-connectivity is different between correct/wrong and baseline conditions. Thin/thick line represents statistical significance at the p-value level of 0.05/0.001. Statistical significance between conditions measured by Fisher’s z-transformation. (C) Heatmap of OB-mPFC power correlation between all pairs of studied frequencies in the baseline (leftmost panel), wrong (middle panel), and correct (rightmost panel) conditions. Solid vertical and horizontal black lines denote borders of frequency bands. (D) Heatmap of OB-mPFC power correlation’s statistical significance, measured by Fisher’s z-transformation, each pair of conditions. Solid vertical and horizontal black lines denote borders of frequency bands. OB, olfactory bulb; mPFC, medial prefrontal cortex.

Next, we computed coherence, which is a neural signature of emotional and cognitive processes, such as WM^9-11,16-18,33,45,47^. As shown in Fig. 5A, WM synchronizes the OB-mPFC circuit, regardless of animal’s performance (statistical significance denoted in horizontal green and pink lines and should be interpreted as in Fig. 2A). Additionally, theta coherence is greater in correct trials, compared to wrong trials (mean ± SEM; coherence_correct_ = 0.50 ± 0.07, coherence_wrong_ = 0.34 ± 0.05, p = 0.06; Fig. 5A,B); however, this effect, despite presence of the trend, did not reach significance in delta (mean ± SEM; coherence_correct_ = 0.15 ± 0.09, coherence_wrong_ = −0.05 ± 0.08, p = 0.10; Fig. 5A,B) and beta (mean ± SEM; coherence_correct_ = 0.24 ± 0.05, coherence_wrong_ = 0.13 ± 0.04, p = 0.11; Fig. 5A,B) bands. Our evaluation of effect sizes showed that, when compared to baseline connectivity, correct trials had weak to moderate (average d’_delta_ = 0.24, 0.13 < d’_delta_ < 0.33) and moderate to large (average d’_beta_ = 0.52, 0.24 < d’_beta_ < 0.92) enhancements in delta and in beta, respectively, along with a strong theta (average d’_theta_ = 1.00, 0.37 < d’_theta_ < 1.26) synchronization (green trace in Fig. 5C); in this case, wrong trials had negligible magnitude in delta (average d’_delta_ = −0.10, −0.17 < d’_delta_ < −0.03) oscillations, accompanied by large theta (average d’_theta_ = 0.77, −0.02 < d’_theta_ < 1.16) and moderate beta (average d’_beta_ = 0.31, −0.04 < d’_beta_ < 0.97) effects (red trace in Fig. 5C). Direct comparisons of correct and wrong trials showed minor differences in delta (average d’_delta_ = 0.26, 0.18 < d’_delta_ < 0.34), low theta (average d’_low theta_ = 0.34, 0.18 < d’_low theta_ < 0.41), and beta (average d’_beta_ = 0.20, −0.02 < d’_beta_ < 0.38; especially in mid-beta with average d’_mid-beta_ = 0.24, 0.10 < d’_mid-beta_ < 0.32) bands (blue trace in Fig. 5C). Further, we found that the animals’ performance is correlated with coherence in high delta, low theta, and mid-beta (grey trace in Fig. 5C); interestingly, the frequencies whose coherence was correlated with behavior were generally associated with larger correct-wrong d’ (compare blue and grey traces in Fig. 5C). Together, it seems that the OB-mPFC phase-based connectivity predominantly happens through theta oscillations. Notably, power-based connectivity in this circuit was stronger in delta and beta oscillations (see Fig. 4).

**Figure 5.**
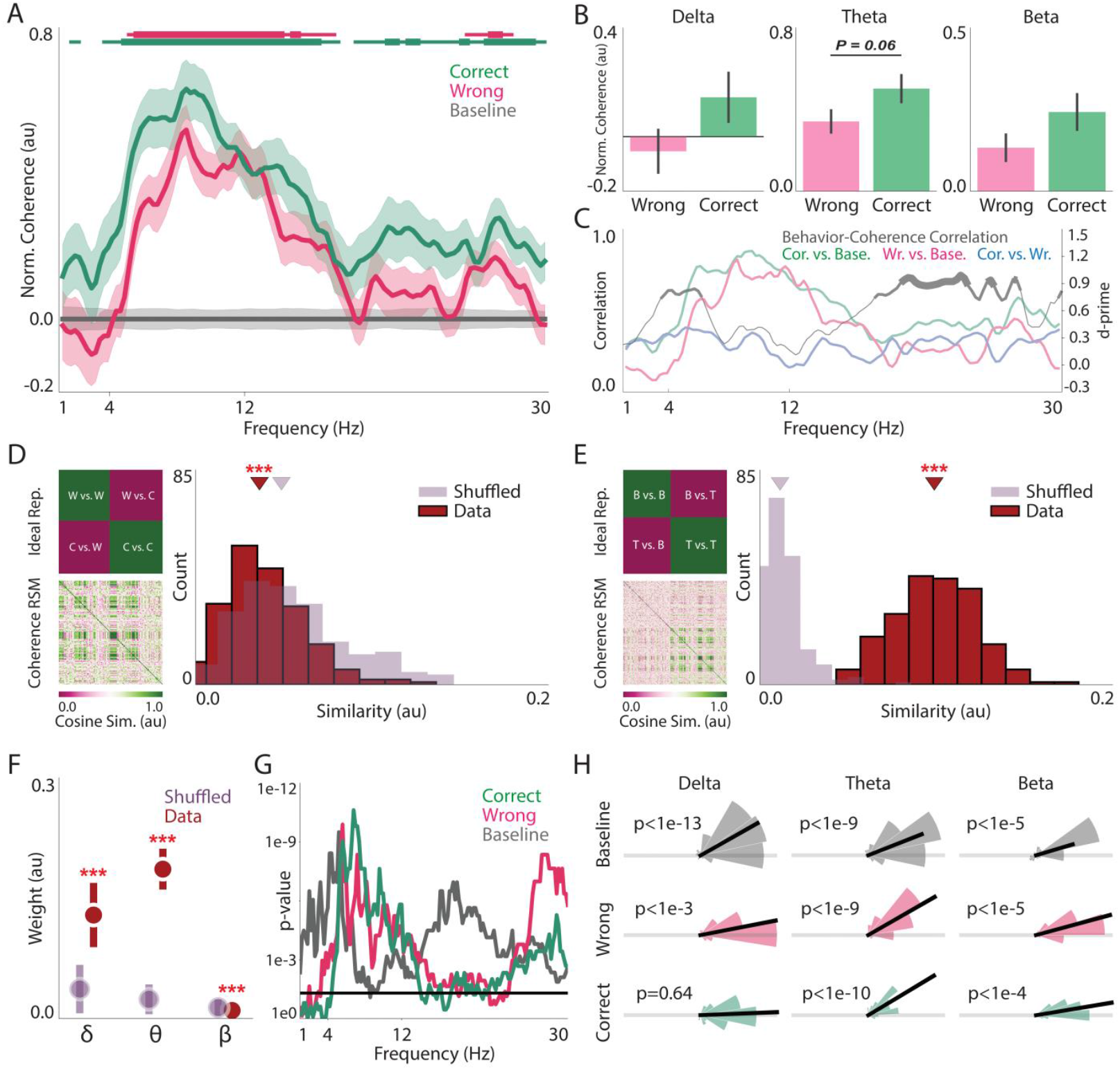
OB-mPFC slow oscillation phase-based connectivity during WM. (A) Normalized OB-mPFC slow oscillations spectral coherence during baseline and task. Solid line and shaded area denote mean and SEM of coherence, respectively. Horizontal green/pink line indicates frequencies with statistical difference between correct/wrong and baseline trials. Thin/thick line represents statistical significance at the p-value level of 0.05/0.001. Permutation test. (B) Statistical comparison of average delta (leftmost panels), theta (middle panels), and beta (rightmost panels) OB-mPFC coherence between correct and wrong trials. Bars and error bars represent mean and SEM, respectively. Permutation test. (C) Frequency-wise coherence-behavior correlation (left y-axis) and magnitude of coherence difference, i.e., d’, between conditions (right y-axis). Thin correlation line indicates values with p ≥ 0.1, while medium/thick line indicates correlation values with p < 0.1/0.05. (D,E, left panels) Ideal (upper panels) and coherence-derived (upper panels) RSMs for correct-wrong (D) and task-baseline (E) conditions. (D,E, right panels) Histogram of the similarity between coherence-derived and ideal RSMs for correct-wrong (D) and task-baseline (E) conditions. Permutation test. (F) Statistical comparison of the classifiers’ weights, trained on coherence data for task vs. baseline, between normal and label-shuffled classification for each frequency band. Permutation test. Point and error bars represent mean and SD, respectively. (G) Frequency-wise p-value for the difference of OB-mPFC phase-lag from 0 rad. Solid horizontal black line indicates p = 0.05. Binomial test. (H) Same as (G), but for average OB-mPFC phase-lag in delta (leftmost column), theta (middle column), and beta (rightmost column) bands. *** p < 0.001. OB, olfactory bulb; mPFC, medial prefrontal cortex; RSM, representational similarity matrix.

We wondered whether the pattern of OB-mPFC coherence can represent WM-related information. Using RSA, we found that while slow oscillations coherence cannot form correct-wrong representation (mean ± SD; similarity_observation_ = 0.009 ± 0.006, similarity_shuffled_ = 0.012 ± 0.008, p < 1e-4; permutation test, Fig. 5D), they convey information on task-baseline distinction (mean ± SD; similarity_observation_ = 0.10 ± 0.02, similarity_shuffled_ = 0.01 ± 0.01, d’ = 4.56, p < 1e-4; permutation test, Fig. 5E). This latter categorization, i.e., task vs. baseline, was also achieved by an LDA classifier (mean ± SD; accuracy_observation_ = 75.00 ± 4.11, accuracy_shuffled_ = 50.32 ± 5.53, p < 1e-4; permutation test; data not shown), whose weights were then extracted and used to define the contribution of different frequency bands (as in Fig. 3E); we observed that delta (mean ± SD; weight_observation_ = 0.13 ± 0.04, weight_shuffled_ = 0.04 ± 0.03, p < 1e-4; permutation test; Fig. 5F) and theta (mean ± SD; weight_observation_ = 0.19 ± 0.03, weight_shuffled_ = 0.02 ± 0.02, p < 1e-4; permutation test; Fig. 5F), but not beta (mean ± SD; weight_observation_ = 0.01 ± 0.01, weight_shuffled_ = 0.01 ± 0.01, p = 1e-4; permutation test; Fig. 5F), weights were greater in the data, compared to shuffled classification, with the theta oscillations showing the most prominent contributions (d’_delta_ = 2.60, d’_theta_ = 7.27, d’_beta_ = −0.38). Also, similar to power (Supplementary Fig. 1B), baseline coherence does not form task-informative representations (mean ± SD; similarity_observation_ = 0.16 ± 0.03, similarity_shuffled_ = 0.01 ± 0.02, d’ = 5.38, p < 1e-4; permutation test, Supplementary Fig. 1A,C). Overall, the pattern of OB-mPFC circuit coherence encodes task-related information, with theta oscillations playing the most crucial role in this process.

Besides synaptic information transfer, which is the typical form of inter-regional communications between brain structures, volume-conduction can also lead to detection of enhanced functional connectivity^57^. Thus, we tried to see whether the connectivity results observed here are synaptically-conducted, by ruling-out the possibility of volume conduction. If there is volume conduction between two neural signals, they will have a phase-lag of zero or π^57^. In the current data, we found that in most cases the phase-lag between OB and mPFC oscillations differs from zero and π, except for delta oscillations during correct trials (statistical significance measured with binomial test; statistics are provided within Fig. 5G,H). Therefore, overall, we show that delta and beta power and theta phase mediate the functional connectivity between OB and mPFC during WM, most likely through synaptic processes.

### mPFC-to-OB slow band information transfer during WM

Up to now, we observed power- and phase-based connectivity between OB and mPFC during WM execution. Subsequently, we sought to find the information flow, if any, between OB and mPFC, using Granger causality (GC). We observed broad band signal transduction in mPFC-to-OB direction in correct and wrong trials (Fig. 6A, left panel; statistical significance denoted in horizontal green and pink lines and should be interpreted as in Fig. 2A). When compared to baseline, both conditions had medium to large effects in nearly all frequencies, with correct trials being relatively larger (correct: average d’ = 0.67, 0.43 < d’ < 0.87; wrong: average d’ = 0.53, 0.42 < d’ < 0.63; green and red traces in Fig. 6B, left panel, respectively); also, there was a moderate difference between correct and wrong settings (average d’ = 0.30, 0.21 < d’ < 0.38; blue trace in Fig. 6B, left panel). Moreover, theta as well as low- and mid-beta GC was correlated with animals’ behavioral performance (grey trace in Fig. 6B, left panel). OB-to-mPFC directionality existed in the theta and beta, but not delta, bands (Fig. 6A, right panel; statistical significance denoted in horizontal green and pink lines and should be interpreted as in Fig. 2A). When compared to baseline, both trial types had medium effects in theta (correct: average d’ = 0.59, 0.28 < d’ < 0.71; wrong: average d’ = 0.53, 0.34 < d’ < 0.70; green and red traces in Fig. 6B, right panel, respectively) and beta (correct: average d’ = 0.43, 0.35 < d’ < 0.50; wrong: average d’ = 0.29, 0.20 < d’ < 0.42; green and red traces in Fig. 6B, right panel, respectively) bands, with theta GC showing the largest magnitudes in both cases and correct trials being slightly higher. The correct-wrong difference was negligible nearly everywhere (average d’ = 0.00, −0.17 < d’ < 0.20; blue trace in Fig. 6B, right panel) and this directionality was not correlated with behavior (grey trace in Fig. 6B, right panel).

**Figure 6.**
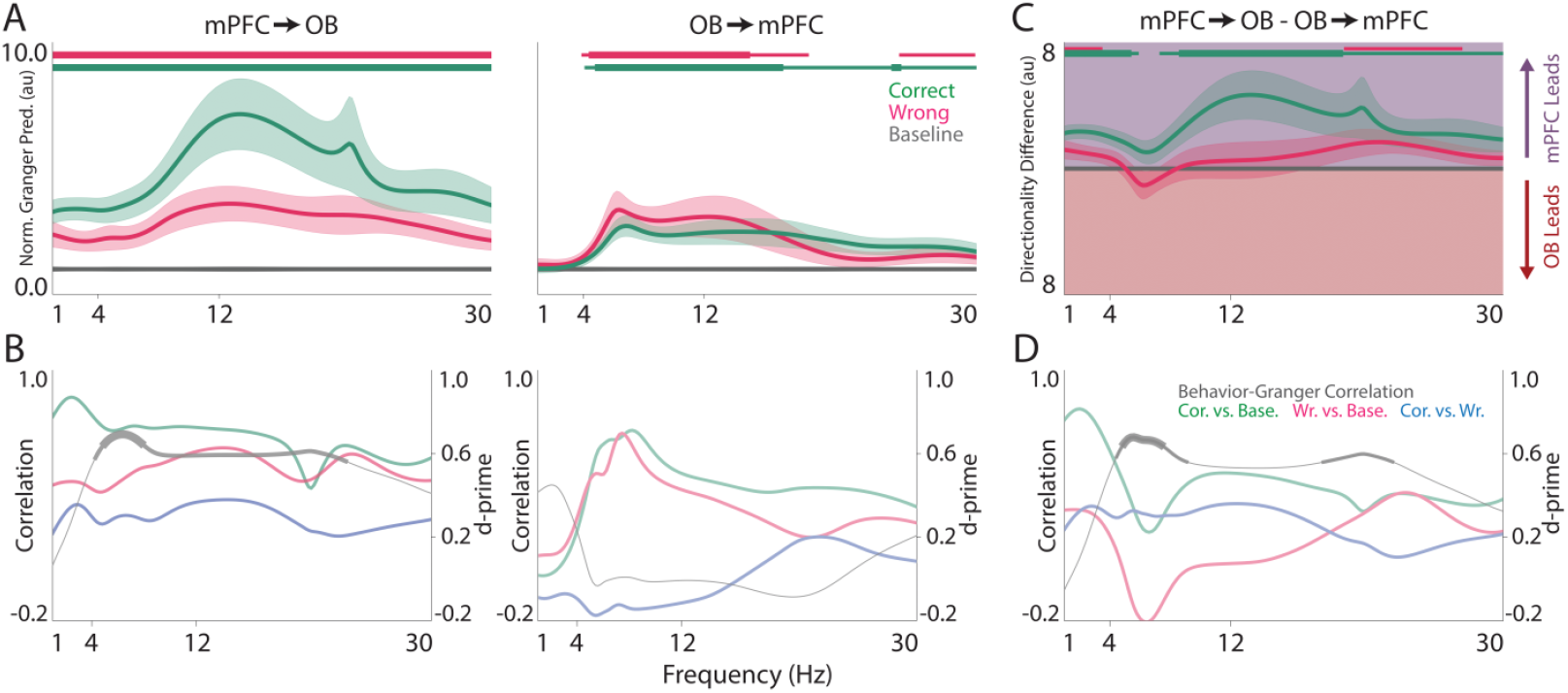
Information transfer between OB and mPFC during WM. (A,C) Normalized GC in the mPFC-to-OB (A, left panel) and OB-to-mPFC (A, right panel) directions as well as directionality difference (C) in baseline, correct, and wrong conditions. Solid line and shaded area denote mean and SEM, respectively. Horizontal green/pink line indicates frequencies with statistical difference between correct/wrong and baseline trials. Thin/thick line represents statistical significance at the p-value level of 0.05/0.001. Permutation test. (B) Frequency-wise GC-behavior correlation (left y-axes) and magnitude of GC difference i.e., d’, between conditions (right y-axes) mPFC-to-OB (left panel) and OB-to-mPFC (right panel). Thin correlation line indicates values with p ≥ 0.1, while medium/thick line indicates correlation values with p < 0.1/0.05. OB, olfactory bulb; mPFC, medial prefrontal cortex

The net OB-mPFC circuit information transfer, computed as the difference of the two directions, showed that mPFC leads OB during correct trials in all frequencies, except for low theta band (Fig. 6C; statistical significance denoted in horizontal green line and should be interpreted as in Fig. 2A). Also, nearly all frequencies showed moderate effects (average d’ = 0.45, 0.22 < d’ < 0.82), with delta (average d’ = 0.76, 0.61 < d’ < 0.82) and low theta (average d’ = 0.35, 0.22 < d’ < 0.58) oscillations being the largest and smallest, respectively (Fig. 6D, green trace). Delta and high-beta conveyed mPFC signals to OB during wrong trials (Fig. 6C; statistical significance denoted in horizontal pink line and should be interpreted as in Fig. 2A), and had moderate effects, in comparison to baseline (delta: average d’ = 0.30, 0.21 < d’ < 0.33; high-beta: average d’ = 0.29, 0.22 < d’ < 0.41; red trace in Fig. 6D); other frequencies had negligible magnitude of difference. Furthermore, delta (average d’ = 0.32, 0.24 < d’ < 0.35), theta (average d’ = 0.32, 0.29 < d’ < 0.36), and low-beta (average d’ = 0.32, 0.25 < d’ < 0.36) bands had moderate correct-wrong d’ (blue trace in Fig. 6D). Also, theta and mid-beta net directionality was correlated with behavior (grey trace in Fig. 6D). Overall, these results imply that there is bidirectional information transfer between mPFC, as a higher order cognitive region, and OB, as a sensory gateway to the brain, during a WM process.

### Delta and theta phase modulate beta amplitude during WM

Visual WM in primates is associated with delta^12^ and theta^8^ - beta phase-amplitude coupling (PAC) in PFC. Therefore, we sought to figure out whether delta and theta phase can modulate regional beta activity. In mPFC, delta (mean ± SEM; PAC_baseline_ = 1.00 ± 0.03, PAC_correct_ = 1.68 ± 0.17, p_correct-baseline_ < 1e-4; PAC_wrong_ = 1.74 ± 0.22, p_wrong-baseline_ < 1e-4; permutation test; Fig. 7A,B and Fig. 7C, left panel) and theta (mean ± SEM; PAC_baseline_ = 1.00 ± 0.03, PAC_correct_ = 2.25 ± 0.20, p_correct-baseline_ < 1e-4; PAC_wrong_ = 2.00 ± 0.17, p_wrong-baseline_ < 1e-4; permutation test; Fig. 7A,B and Fig. 7C, right panel) modulation of regional beta activity was augmented during both correct and wrong trials, compared to baseline. We also observed similar effects in the OB (mean ± SEM; delta: PAC_baseline_ = 1.00 ± 0.03, PAC_correct_ = 1.62 ± 0.15, p_correct-baseline_ < 1e-4; PAC_wrong_ = 1.45 ± 0.16, p_wrong-baseline_ = 0.0004; theta: PAC_baseline_ = 1.00 ± 0.03, PAC_correct_ = 3.14 ± 0.24, p_correct-baseline_ < 1e-4; PAC_wrong_ = 3.12 ± 0.23, p_wrong-baseline_ < 1e-4; permutation test; Fig. 7D-F). There was no significant regional delta- or theta-beta PAC difference between correct and wrong conditions, in neither region (all cases had a p > 0.05; Fig. 7A-F).

**Figure 7.**
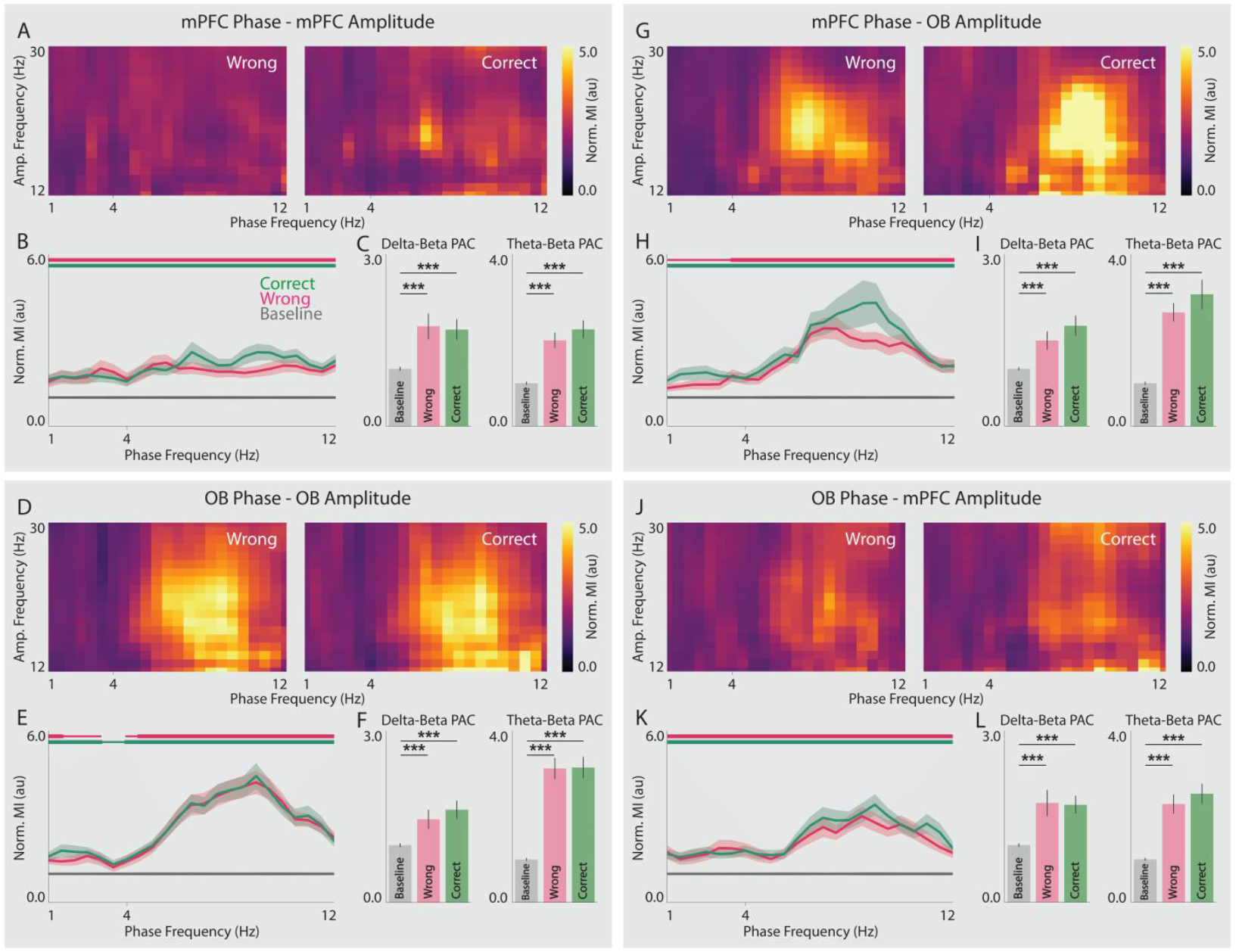
Local and long-range delta- and theta-beta PAC between OB and mPFC during WM. (A,D,G,J) Average of normalized MI in correct (right panels) and wrong (left panels) trials for mPFC-mPFC (A), OB-OB (D), mPFC-OB (A), and OB-mPFC (D). (B,E,H,K) Normalized MI of mPFC-mPFC (B), OB-OB (E), mPFC-OB (H), and OB-mPFC (K) PAC during baseline, wrong, and correct trials. Solid line and shaded area denote mean and SEM, respectively. Horizontal green/pink line indicates frequencies with statistical difference between correct/wrong and baseline trials. Thin/thick line represents statistical significance at the p-value level of 0.05/0.001. (C,F,I,L) Statistical comparison of average delta-(left panels) and theta-(right panels) beta PAC between baseline, wrong, and correct trials. Bars and error bars represent mean and SEM, respectively. Permutation test. ***p < 0.001. OB, olfactory bulb; mPFC, medial prefrontal cortex; MI, modulation index; PAC, phase-amplitude coupling.

We next tried to figure out whether delta and theta oscillations of either region can modulate the beta activity in the other area. We found that PFC delta (mean ± SEM; PAC_baseline_ = 1.00 ± 0.02, PAC_correct_ = 1.75 ± 0.17, p_correct-baseline_ < 1e-4; PAC_wrong_ = 1.49 ± 0.15, p_wrong-baseline_ < 1e-4; permutation test; Fig. 7G,H and Fig. 7I, left panel) and theta (mean ± SEM; PAC_baseline_ = 1.00 ± 0.03, PAC_correct_ = 3.07 ± 0.33, p_correct-baseline_ < 1e-4; PAC_wrong_ = 2.65 ± 0.20, p_wrong-baseline_ < 1e-4; permutation test; Fig. 7G,H and Fig. 7I, right panel) oscillations modulated OB beta activity during correct and wrong trials. On the other hand, mPFC beta band was tuned by OB delta (mean ± SEM; PAC_baseline_ = 1.00 ± 0.02, PAC_correct_ = 1.70 ± 0.15, p_correct-baseline_ < 1e-4; PAC_wrong_ = 1.74 ± 0.22, p_wrong-baseline_ < 1e-4; permutation test; Fig. 7J,K and Fig. 7L, left panel) and theta (mean ± SEM; PAC_baseline_ = 1.00 ± 0.03, PAC_correct_ = 2.53 ± 0.23, p_correct-baseline_ < 1e-4; PAC_wrong_ = 2.29 ± 0.21, p_wrong-baseline_ < 1e-4; permutation test; Fig. 7J,K and Fig. 7L, left panel) oscillations during both trial types. We observed no significant differences between correct and wrong trials in the inter-regional PAC within the OB-mPFC circuit (all cases had a p > 0.05; Fig. 7G-L). Together, we suggest that these inter-regional delta- and theta-beta PAC in the OB-mPFC circuit support distinct neural processes. OB, as a global coordinator of neural activity^11,14,17,18,23,34^, can potentially mediate respiratory signals to modulate the activity of mPFC, possibly other regions, and behavior. In the same manner, mPFC can exert top-down cognitive control and filter task-relevant information from the brain input by affecting OB activity.

## Discussion

Here, we showed that the dynamics of OB-mPFC slow oscillations support rodent WM. Regional delta, theta, and beta activities are elevated during WM, and interestingly the pattern of these activities can distinguish whether or not the animal is involved in a cognitive process. We also found that these regions cooperate to support WM with a variety of mechanisms: 1) enhanced power- and phase-based functional connectivity; 2) bidirectional information transfer; and 3) tuning of beta activity by regional and inter-regional delta and theta phase.

A large body of literature focuses on how the neural activity in a given brain region supports a specific mammalian behavior^9,13-19,25,29,32,45,47^. In case of WM, PFC is thought to be the region with the most prominent roles, acting through diverse mechanisms^1,3,4,8-12,20-22,24,25,27-29,31-33^. Systemic and/or neuropsychiatric disorders that alter mPFC activity cause malfunctions in WM^9,32^. On the other hand, while faster neural activities, such as gamma oscillations, are thought to convey sensory input to the nervous system, slow rhythms are associated with more cognitively advanced processes, such as WM, and are suggested to mediate the cognitive control exerted by higher order areas to manage global brain resources^3,21,22^. Our results are in line with both perspectives; specifically, slow band activities in mPFC, and more interestingly OB, are enhanced while the animals are engaged in a WM process; in many frequencies, these enhancements are greater during correct, compared to wrong, trials and are corelated with the subjects’ task performance, further signifying their behavioral relevance. Moreover, there is extensive evidence on enhancement of OB activity and/or its modulations over other brain regions during behavior^10,11,14,17,18,33-35,58^. Mechanistically, OB oscillations are largely driven by mechanical stimulation of olfactory sensory neurons (OSNs) by respiration^17,59-61^, whose improper functioning directly disrupts animal behavior^17^. Notably, such behavioral abnormalities are also observed following perturbation of OSNs or OB activity through a range of methods^14,17,32^, and electrical stimulation of either can positively affect the function of mPFC and OB and improve memory^32,58^. Thus, slow band activities of mPFC and OB have the potential to regulate mammalian cognitive performance and/or emotional/psychiatric status.

Synchronization across different brain regions supports diverse behaviors^3,9-11,13-19,24,25,32,33,45,47,48^. Both PFC and OB are functionally connected to other structures and integrated in networks that substantially shape sensory and/or cognitive functions, emotions, and even circuit developments^4,9-11,13-19,22,24,25,32,33,36,45,62^. Here, we showed a variety of coupling mechanisms in the OB-mPFC circuit during WM. Specifically, we observed enhanced delta and beta power connectivity (see Fig. 4). Conversely, while phase coupling was broad band, it was more prominent in theta oscillations (see Fig. 5). Importantly, the connectivity enhancements were generally larger in correct, compared to wrong, trials. Slow band coupling in this circuit have been linked to several behaviors, however with different pattens^10,14,17-19,32^. For instance, as we have previously shown, OB-mPFC theta power connectivity accompanies rodent anxiety^18^; here however, we did not find this type of communication during WM. We also observed that slow oscillations are a route for information transfer between the two structures during WM (see Fig. 6). Further, oscillations in either region tunes the activity of the other area (see Fig. 7). These bidirectional neural interactions are well-aligned with the feed-forward and feedback processes expected from hierarchically-organized circuits and networks in the brain^1^. In this view, while the upstream, often more primitive region, conveys task-related information to higher order structures, the downstream, generally more cognitively advanced area, controls the brain input by exerting feedback modulations^1^. Interestingly, the roles of such interactions are highlighted during WM/attention^1^.

In contrast to single neuron processing perspectives, the pattern of activity/activations in a population of neurons is a rich repertoire of neural codes that govern behaviors^52-56^. Here, we proposed a similar hypothetical neural space, in which dimensions are defined by the oscillatory activity/connectivity in a given brain region/circuit at different frequencies (see Fig. 3A, right panel). Next, to provide experimental evidence for this framework, we showed that the pattern of OB and mPFC local activity as well as their functional connectivity can detect the animal’s experience of a cognitive process. Often times, behaviors are sticked to specific lower dimensional subspaces, i.e., neural manifolds, which is the small number of all hypothetically possible activity patterns that define the entire space^53-56^. Unlike single neuronal activities that are subject to change over time, the latent space representations derived from neuronal activity patterns are shown to be stable in time^63^ and even transferrable across individuals^64^. Since population activity is a strong predictor of neural oscillation^65^, presence of such stable manifolds in the oscillatory-based neural space is not unexpected. Therefore, successful completion of behaviors might be associated with specific neural trajectories in these spaces.

Together, our results propose the following frameworks: 1) the dynamics of mPFC and OB neural activities can influence animal behavior through local or long-range interactions, with implications that extent beyond the scope of current study findings; 2) patterns of activity/connectivity across brain regions/circuits encode task-related information and high-dimensional neural spaces formed by brain oscillations may contain representations or manifolds with behavioral correspondence. This perspective could open new avenues for uncovering the functional significance of brain rhythms. We encourage future investigations to test these hypotheses across diverse behavioral contexts and to employ neural perturbation approaches for establishing causal relationships.

## Methods

### Animals

Ten healthy male Wistar rats, aged 8-10 weeks and weighing 200-250 gr, were obtained from the Laboratory Animals Research Center at Zahedan University of Medical Sciences (Zahedan, Iran). The animals were housed under standard condition, temperature at 21 ± 2 °C with 12-hour light/dark cycle, and free access to water and food throughout the study. All procedures were performed consistent with the National Research Council’s Guide for the Care and Use of Laboratory Animals and the ARRIVE 2.0 guidelines. Study protocols and procedures were approved by the Ethics Committee of Zahedan University of Medical Sciences (IR.ZAUMS.AEC.1404.001).

### Surgery

Surgical procedures were similar to our previous study^18^. Briefly, general anesthesia was inducted by intraperitoneal injection of ketamine (100 mg/kg) and xylazine (10 mg/kg). To maintain the body temperature at 37 °C, a heating pad placed beneath the animal’s body. Appropriate anesthetic depth confirmed by lack of pinch and tail reflexes. Then, animals placed in a stereotaxic frame (Narishige, Japan) and received a subcutaneous injection of lidocaine (0.5 ml) used for local anesthesia at the site of surgery. Two stainless-steel recording electrodes (127 µm diameter; A.M. Systems Inc., USA) were implanted into the left OB (coordinates: AP −8.5 mm, ML −1 mm, DV −1.5 mm) and prelimbic PFC (coordinates: AP +3.2 mm, ML −0.6 mm, DV −3.6 mm). A stainless-steel screw was inserted into right parietal bone as the reference electrode. All electrodes were connected to a socket that stabilized using dental acrylic cement. Following the procedure, the surgical site was disinfected using tetracycline ointment. Post-operative care was performed with buprenorphine injection (0.1 mg/kg) for analgesia and sterile saline injection (1.0 ml) for hydration. Afterwards, the animal was returned to their home cages for recovery.

### Electrophysiological recordings

After 7 days of recovery, simultaneous LFPs were recorded in real time from OB and mPFC while the animals performed the behavioral test. Throughout the recordings, head-mounted socket was connected to a miniature, high-input impedance buffer head-stage (BIODAC-A, TRITA Health Tec. Co., Tehran, Iran), that was linked via cables to a primary AC-coupled amplifier (gain: ×1000) and the data acquisition system (BIODAC-Bi401l9B, TRITA Health Tec. Co., Tehran, Iran). LFPs from both OB and mPFC were recorded simultaneously at 2 kHz sampling rate (with a low-pass filter of 250 Hz) stored for further offline analyses. All signal processing and data analyses were performed using Matlab 2019b (The MathWorks Inc., USA) and Python v3.8.20.

### Behavioral procedure

To reduce animal’s stress during test, all animals were habituated to the recording environment one week prior and following the surgery. The recording sessions consisted of two separate sections. At the first section, to achieve baseline OB and mPFC activity, animals were placed in their home cage and simultaneous LFP were recorded for 10 min. Next, the recording was made while the animals explored the Y-maze apparatus. Y-maze consisted of three identical arms (50 cm in length, 10 cm in width, and 25 cm-high walls) arranged at 120° with respect to each other. The animals were allowed to freely explore the test environment for 10 min. Sequential transitions that contained visiting all arms consecutively were considered as correct trials (Fig. 1B). A video camera recorded the animals’ behavior in the during the tasks.

### Histological verification of electrode placements

Following the completion of the recording sessions, animals were deeply anesthetized with an intraperitoneal injection of urethane (1.2 g/kg). To mark the electrode tips, small electrical lesions were made before the brain extraction. Then, the brains were removed and fixated in 4% paraformaldehyde for 48 hours. Coronal brain sections (200 μm thick) were prepared and examined by light microscopy to verify electrode placements within the OB and mPFC.

### Neural data analyses

We considered 5-sec moving periods of home cage exploration and every transition in the middle triangle of Y-maze (longer than 1 sec) as baseline (n = 120) and task (n_correct_ = 73, n_wrong_ = 70) trials, respectively. PSD of the LFP signals was estimated using custom written Matlab codes. A complex Morlet wavelet was convolved with the signal to obtain time-frequency representations across frequency range (1-30 Hz, step of 0.2 Hz). Average power values across time were computed for each trial as the raw power for subsequent analyses. To normalize power values, the mean and standard deviation of baseline trials’ power were calculated separately for each animal. Then, power values of each trial were z-scored to this baseline activity at each frequency.

To assess to OB-mPFC functional connectivity, power-correlation between OB and mPFC activity was calculated by spearman’s rank correlation in Python (SciPy ***spearmanr*** function) separately for correct, wrong, and baseline trials. We also computed magnitude-squared coherence between OB and PFC signals in different conditions using chronux toolbox^66,67^ in Matlab. Raw coherence values were normalized to baseline connectivity similar to PSD analysis.

We investigated the directionality between OB and mPFC using GC^68,69^, as suggested earlier^18,57^. For that, we down-sampled LFP signals from 2kHz to 500 HZ to enhance stability of GC autoregressive (AR) models and improve signal to noise ratio. We set the AR model order to 10 and computed GC for both OB-to-mPFC and mPFC-to-OB directions. For normalization, GC values are reported proportional to the baseline GC in each direction separately.

To compute the phase-lag between mPFC and OB signals, we first extracted the phase values of each region by finding the angle of the analytical signal of the wavelet transform. Next, phase differences were calculated by subtracting the instantaneous phases of the two regions, which were then averaged across time. These values were statistically compared to 0 and π.

To compute PAC, LFP phase values were extracted using Hilbert transform and were divided into 18 phase bins (20° each). Also, the average signal amplitude, computed using furrier transform, was calculated in each phase bin. Subsequently, PAC was defined as the magnitude of the mean vector that derived from the phase-binned amplitude values. This vector length counted as the modulation index for subsequent statistical comparisons. For normalization, PAC values are reported proportional to the baseline PAC in each case separately.

To explore the pattern of OB and mPFC activities, we trained LDA classifiers (using scikit-learn ***LinearDiscriminantAnalysis*** function) to either detect correct vs. wrong or task vs. baseline. These procedures were repeated 200 times an in every iteration %70/%30 of the trials was used to train/test the model. Also, to find the distribution of chance performance, the same procedure was repeated for 200 times with every time shuffling the trial labels. Of note, the weights of these models were extracted for further analyses to describe the neural architecture in each case.

We used RSA^52^ to study the structure of representation in regional activity and functional connectivity. For that, we first formed the ideal representational matrix for each case, which is theoretically 0 (or 1) for no (or perfect) similarity. To find the data-derived RSMs, we computed cosine-similarity between every pair of trials in each case (using scikit-learn ***cosine_similarity*** function), as the following:

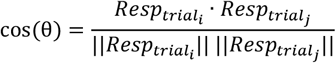

in which 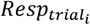 and 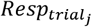 are the vectors of neural response to *trial*_*i*_ and *trial*_*j*_, respectively, and θ is the angle between these two vectors in the high-dimensional neural space. Thus, a greater cos(θ) value suggests more similarity between the two vectors in the presumed space. This process creates the *N* × *N* correct-wrong or task-baseline RSMs with the following structure:

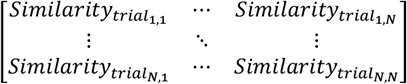

where *N* is total number of trials in each case and 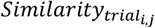is the cosine-similarity between vectors of neural response to *trial*_*i*_ and *trial*_*j*_. Next, the correlation between each data-derived RSM and the ideal representations was calculated with Kendall’s tau correlation (using SciPy ***kendalltau*** function). This procedure was repeated 200 times and, in every run, %70 of the trials was used to form data-derived correct-wrong or task-baseline RSMs. Also, to find the distribution of chance similarities, the same procedure was repeated for 200 times with every time shuffling the trial labels.

### Statistical analyses

Statistical and machine learning analyses were performed in Python v3.8.20, using scikit-learn v1.3.0 and SciPy v1.10.1 libraries, or Matlab 2019b. Circular data statistics were performed with CircStat toolbox^70^ in Matlab. Details of the statistical tests are described at the appropriate context throughout the manuscript. All permutations were repeated 10001 times. Pearson correlations between the animals’ behavioral performance and oscillatory features were done with Matlab ***corr*** function. Correlation values were compared with Fisher’s z-transformation by custom-written Python code. All tests were two-tailed and p-values less than 0.05 were considered as statistically significant. For each pair of data distributions, d’ was computed as:

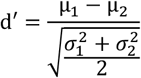

in which μ_*i*_ and σ_*i*_ are the mean and the SD of each distribution, respectively.

## Author Contributions

MM conceptualized the study. ASM and MM performed the surgeries, ran the experiments, collected the data, designed the analysis plan, analyzed the data, and drafted the manuscript. All authors reviewed the manuscript. MRR and MAM supervised the study.

## Acknowledgements

The authors would like to thank Farhad Tabasi and Morteza Salimi for their constructive comments on the experimental design and procedures. We also appreciate technical assistance from Soheil Karimi Darmian.

## Data and Code Availability

Data, Matlab codes, and Python notebooks will be made available upon reasonable request to the corresponding authors.

## Funding

This work was funded by Zahedan University of Medical Sciences (grant number: 11707).

## Competing Interests

None declared.

## Supplementary Information

**Supplementary Figure 1.**
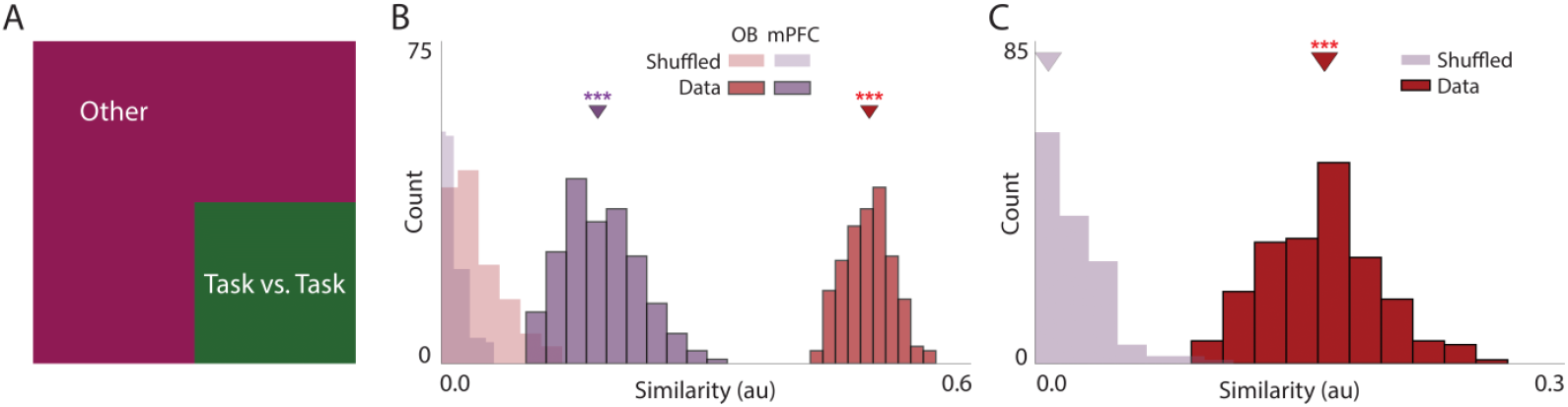
RSA with a cognitive kernel. (A) Ideal task-baseline RSM, under the assumption that the neural activity during baseline is spontaneous and does not represent neuronal processing. (B,C) Histogram of the similarity between power (B) and coherence (C) - derived and ideal RSMs under the assumption described in (A) for the task-baseline setting. Arrowheads denote the median of the similarity distributions. Permutation test. ***p < 0.001. OB, olfactory bulb; mPFC, medial prefrontal cortex; RSA/M, representational similarity analysis/matrix.

## Notes

### Competing Interest Statement

The authors have declared no competing interest.

